# Reward Expectation Reduces Representational Drift in the Hippocampus

**DOI:** 10.1101/2023.12.21.572809

**Authors:** Seetha Krishnan, Mark E.J. Sheffield

**Affiliations:** Department of Neurobiology and Institute for Neuroscience, University of Chicago, Chicago, IL 60637, USA

**Keywords:** Dorsal hippocampus, place cells, reward expectation, representational drift, two-photon microscopy, longitudinal calcium imaging

## Abstract

Spatial memory in the hippocampus involves dynamic neural patterns that change over days, termed representational drift. While drift may aid memory updating, excessive drift could impede retrieval. Memory retrieval is influenced by reward expectation during encoding, so we hypothesized that diminished reward expectation would exacerbate representational drift. We found that high reward expectation limited drift, with CA1 representations on one day gradually re-emerging over successive trials the following day. Conversely, the absence of reward expectation resulted in increased drift, as the gradual re-emergence of the previous day’s representation did not occur. At the single cell level, lowering reward expectation caused an immediate increase in the proportion of place-fields with low trial-to-trial reliability. These place fields were less likely to be reinstated the following day, underlying increased drift in this condition. In conclusion, heightened reward expectation improves memory encoding and retrieval by maintaining reliable place fields that are gradually reinstated across days, thereby minimizing representational drift.

## Introduction

Representational drift refers to the changes in neural representation that occur in the same environment, despite seemingly stable behavioral output (Driscoll et al., 2022; Mau et al., 2020; Rule et al., 2019). This phenomenon has been observed in many brain regions, including sensory and motor areas, as well as in memory areas like the hippocampus (Delamare et al., 2023; Driscoll et al., 2017; Lütcke et al., 2013; Marks & Goard, 2021). In the hippocampus, place cells encode an animal’s position in an environment and are thought to represent spatial memories. Place cells have been shown to drift gradually over the course of several days. Many place cells emerge and vanish from the place code over time, resulting in a reduction of spatial activity correlation across multiple days (Dong et al., 2021; Gonzalez et al., 2019; Hainmueller & Bartos, 2018; Ziv et al., 2013). Yet, drift can also be more drastic, with entire populations of place cells ceasing to fire and a new population emerging (remapping), despite no changes to the environment (Low et al., 2021; Rubin et al., 2015; Sheintuch et al., 2020). Interestingly, the location decoding ability by place cells is generally preserved during drift (Gonzalez et al., 2019; Keinath et al., 2022; Ziv et al., 2013). Drift is thus believed to contribute to how the brain continuously learns, allowing representations to be flexible rather than rigid and stable, yet preserving sufficient information over time to precisely retrieve memories (Driscoll et al., 2022; Mau et al., 2020; Rubin et al., 2015; Taxidis et al., 2020).

Not all memories are created equal. Changes in the internal state of the animal have been suggested as an important contributor to drift (Hulse et al., 2017; Kentros et al., 2004; Niell & Stryker, 2010; Sadeh & Clopath, 2022; Vinck et al., 2015). Internal states, such as the expectation of a reward, influence memory formation and retrieval (Krishnan et al., 2022; Schultz et al., 1997). Indeed, memories of environments where a reward is expected are better remembered compared to those where it is not (Ambrose et al., 2016; Braun et al., 2018; Gruber et al., 2016; Singer & Frank, 2009). These more salient memories are often described as being more stable over time and therefore retrieved more accurately (Goode et al., 2020; Josselyn & Tonegawa, 2020; Lütcke et al., 2013; Pettit, Yap, et al., 2022). But how does the hippocampal representation balance the extent of drift versus stability to differentiate memories of different valences within the same environment? Relatedly, do poorly retrieved memories result from greater drift compared to well retrieved memories?

In a recent study (Krishnan et al., 2022), we found that when reward expectation is lowered in an environment, many place cells encoding that environment undergo remapping and exhibit impoverished spatial encoding properties. This shows that hippocampal representations are dependent on the internal state of the animal. However, an important question remains: do internal states also determine the extent of hippocampal representational drift over time? In this follow-up paper, we set to answer this question. To do so, we employed an experimental paradigm we previously developed (Krishnan et al., 2022) to alter internal states of reward expectation while keeping all other variables constant. Using two-photon calcium imaging, we recorded the activity of the same hippocampal CA1 cells over two days in a familiar rewarded environment with high reward expectation. We then removed the reward to lower reward expectation and recorded the activity of the same cells without reward over two days. We then re-introduced the reward to reinstate reward expectation. This allowed us to directly compare, in the same cells and across days, the extent of drift in hippocampal contextual representations associated with and without reward expectation.

## Results

For ∼2 weeks, mice were trained to run on a treadmill along a 2 m virtual reality (VR) linear track for water rewards delivered at the end of the track (rewarded environment, R1), after which mice were teleported back to the beginning of the track (**Figure 1A**), considered one lap. Well-trained mice exhibited pre-emptive licking for reward, before the rewarded location and before reward delivery, showing evidence of learned reward expectation (Krishnan et al., 2022). These trained mice then underwent a three-day experiment protocol (**Figure 1B**, n = 5 male mice). On Day 1, mice ran in the rewarded environment (R1). On Day 2, mice ran in the same rewarded environment (R2) before water reward was unexpectedly removed (Unrewarded environment: UR1). On Day 3, mice first ran in the unrewarded environment (UR2) before water reward was reintroduced (Re-Rewarded environment: RR). Mice experienced each condition for ten minutes. Over the three days, we imaged from the same dorsal hippocampal CA1 neurons (mean ± 95% confidence intervals, 1376 ± 623 cells in n = 5 mice) expressing the calcium indicator GCaMP6f (Chen et al., 2013), using a two-photon microscope (**Figure 1C**). As reported previously (Krishnan et al., 2022), we used pre-licking as a measure of reward expectation and as before, mice pre-licked for a few laps in UR1 as though still expecting a reward. This pre-licking stopped after a few laps (5.5 ± 3.3 laps) reflecting a transition to a low reward expectation state (**Supplementary Figure 1A-B, 2A**). We defined the laps with pre-licking as those with high reward expectation and the laps after pre-licking stopped as those with low reward expectation. We found that CA1 population activity abruptly changed upon lowered reward expectation, rather than immediately following reward removal, matching our previous results (Krishnan et al., 2022) **(Supplementary Figure 2A-B).**

**Figure 1:**
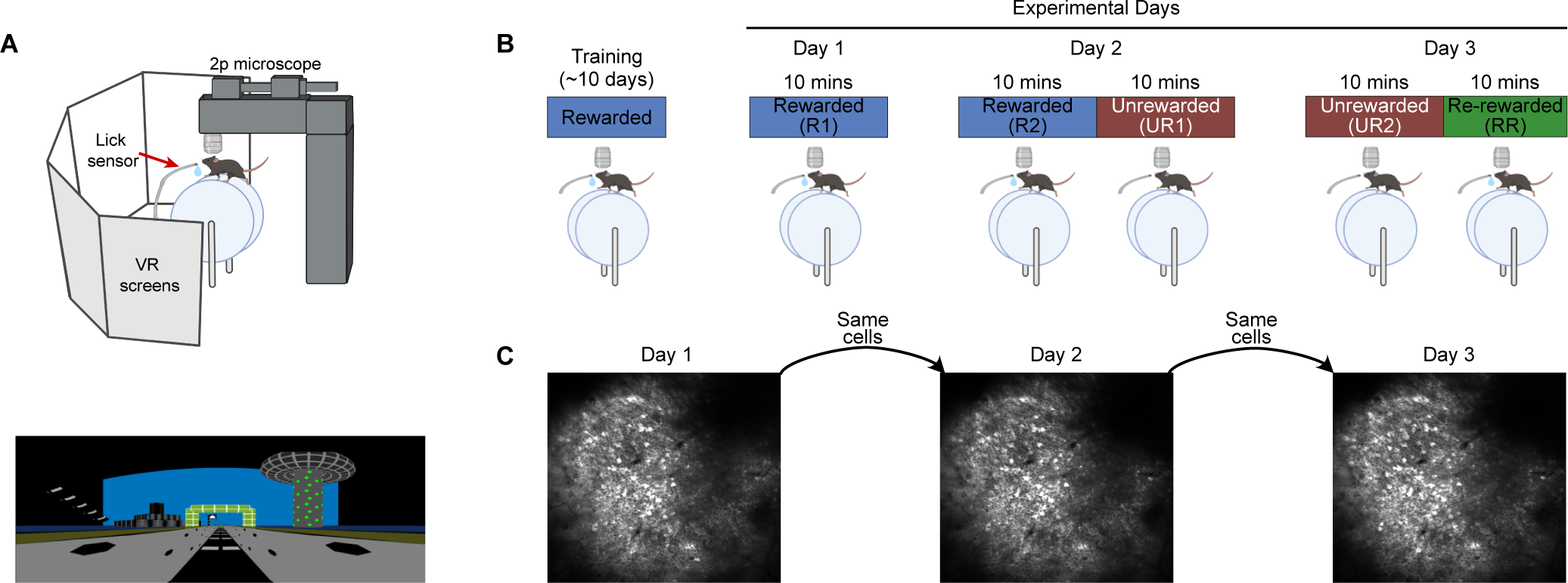
Experimental protocol for tracking place cell dynamics across changing reward expectations and days. (A) (top) Virtual Reality (VR) setup. (bottom) VR environment which is a 2m linear track with several landmarks and visual cues. (B) Three-day experimental protocol, the same VR environment was used across all days and reward contingencies were varied as shown. (C) Example field-of-view obtained from dorsal hippocampus CA1 neurons expressing GCaMP6f. Same field-of-view was aligned and imaged across all three days.

Since this investigation aims to compare the dynamics of activity in the hippocampus across days during high versus low reward expectation states, here onwards, we will only consider the laps in UR1 after mice stopped pre-licking (for a systematic comparison of dynamics between the high and low reward expectation laps within UR1 refer to our previous work (Krishnan et al., 2022)). Notably, we found that this lowered reward expectation state continued in UR2 (**supplementary Figure 1A**). Mice would sometimes lick non-specifically at random locations in UR2 but not consistently across laps (small grey spikes in trace in **Supplementary Figure 1A**). Pre-licking returned a few laps following reward reintroduction in RR as mice re-learned to expect the reward (example in **Supplementary Figure 1B**). Thus, this experimental protocol enabled us to compare the hippocampal dynamics across days between high (R1 and R2) and low (UR1 and UR2) reward expectation states and with the reinstatement of reward expectation (RR).

### Place cells drift more across days when reward expectation is low

We investigated and compared the stability of place fields across days in the two conditions with similar reward expectation states (i.e., R1 to R2 vs UR1 to UR2). Our aim was to test our hypothesis that high reward expectation enhances the stability of place fields across days compared to states of low reward expectation (**Figure 2A-D**). To quantify place field stability, we measured the population vector correlation of all cells between each pair of positions on the virtual track in both conditions (see methods, **Figure 2E, Supplementary Figure 3A)**. We found that CA1 population activity showed greater similarity between R1 and R2 compared to UR1 and UR2 (**Figure 2F-G)** along all parts of the track (**Figure 2G**). This implies that lowering reward expectation increases drift of CA1 spatial representations across days in an unchanging spatial environment.

**Figure 2.**
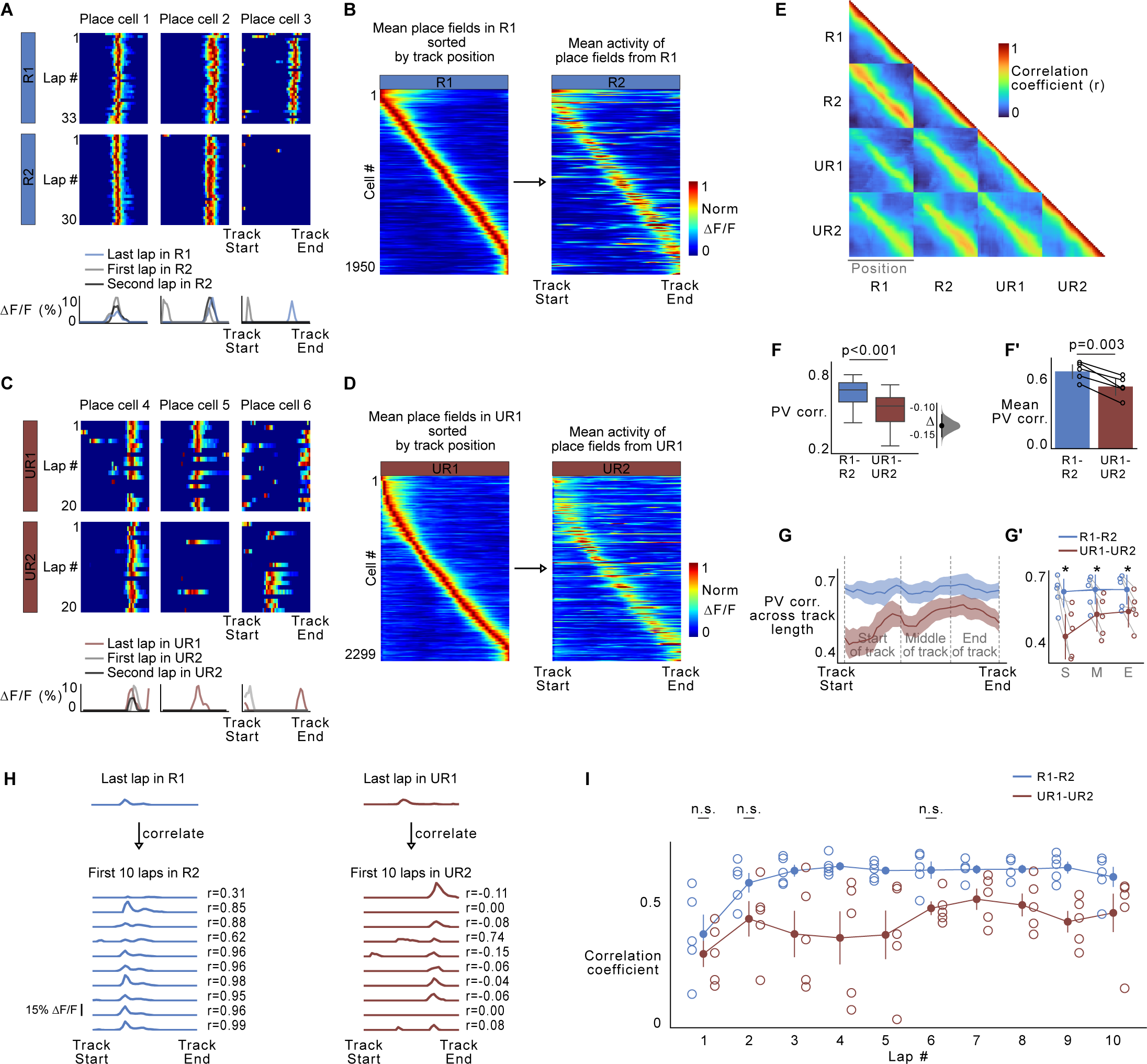
Place cell representations drift more across days when reward expectation is lowered. (A) (top) Example place cells in R1 and R2. (bottom) Traces show activity of the place cell in the last lap in R1 (blue) compared to the first two laps in R2 (grey and black traces). (B) Place fields defined in R1 plotted across R2. Activity of each place cell was normalized to peak in R1 and sorted by its center of mass along the track. (C-D) Same as A-B but between UR1 and UR2. (E) Position vector (PV) correlation between all pairs of position along the track and across all conditions. The correlation was averaged across all imaged cells (n=6880 cells) and all animals (n=5 animals). (F) Distribution of PV correlation between R1-R2 and UR1-UR2 (F’) Animal wise averages of data in F. Each circle represents a single animal. P-value was obtained using a two-sided paired *t* test. (G) PV correlation across track length. Lines represent mean. (G’) Average PV correlation binned by track position indicated by gray lines in the G and averaged across each animal. Circles indicate data from each animal. S: Start of the track, M: Middle of the track, E: End of the track. Asterix (*) denote significant *P* values (two-sided paired *t* test, *P* < 0.01) obtained by comparing R1-R2 correlation (blue) with UR1-UR2 correlation (red) at each position. (H) Examples depicting the lap-by-lap correlation measure quantified in I. The activity of the last lap in R1/UR1 was correlated with activity in the first 10 laps of R2/UR2. Lap traces are shown and are from the same place cell across days. r indicates the resulting correlation coefficient and is listed next to each lap trace. (I) Average correlation coefficient across all place cells defined in R1/UR1 and their correlation with the first 10 laps in R2 (blue) and UR2 (red) respectively. Circles indicate individual mice. *P* values (two-sided paired *t* test, *P* < 0.01) were calculated by comparing R1-R2 correlation (blue) with UR1-UR2 correlation (red) at each lap. n.s. indicates insignificant p-values, all other comparisons were significantly different. Error bars in F’, G’ and I indicate 95% confidence intervals. In G, shaded area represents standard error of the mean (s.e.m). For full details on statistics and p-values, see **supplementary file 1**.

To determine whether the increased drift across UR1 and UR2 was caused by changes in CA1 representations across days rather than within-session changes that may have occurred in UR1 (which could have initially drifted but then stabilized by the end of the session), we examined the correlation of place cell activity on the last lap in UR1 with the first ten laps in UR2. (UR1-UR2, **Figure 2H-I**). We compared this to the correlation of place cell activity on the last lap of R1 with the first ten laps in R2 (R1-R2, **Figure 2H-I**). We found that, on average, correlations between UR1-UR2 remained lower than correlations between R1-R2 (all laps, R1-R2: 0.59 ± 0.06, UR1-UR2: 0.43 ± 0.05, t = 8.26, p<0.001, two-sided paired *t* test, **Figure 2I**). However, when we compared them on a lap-by-lap basis, we found correlations between R1-R2 and UR1-UR2 were not significantly different on the very first lap (R1-R2: 0.39 and UR1-UR2: 0.31, t = 0.895, p = 0.422, two-sided paired *t* test, **Figure 2I**). Interestingly, as the mice ran more laps in R2, the correlation to R1 increased (average second lap correlation with R1: 0.57, and average correlation for laps 2-10: 0.61 ± 0.02). As the mice ran more laps in UR2, the correlation to UR1 increased only modestly and remained lower than when R2 was correlated to R1 (average second lap correlation with UR1: 0.44 vs 0.57 with R1-R2, average correlation for laps 2-10: 0.44 ± 0.04 vs 0.61 with R1-R2, two-sided paired *t* test, **Figure 2I**). This delayed retrieval of the R1 representation in R2 could not be explained by changes in the animal’s running behavior on these initial laps (**Supplementary Figure 3A**). Instead, the previous spatial representation is gradually retrieved over a few laps of the environment, becoming more like the previous day on each lap - dynamics which underlie the low amount of drift in the high reward expectation condition. In the absence of reward expectation, however, gradual retrieval of the previous day’s representation does not occur, explaining the increased drift across days in this condition.

We then investigated how spatial representations change when reward is reinstated (in RR) compared to other conditions. We found that the PV correlation was highest between UR2 and RR compared to R1-RR or R2-RR (**Supplementary Figure 3B-D**). Since it takes a few laps to reinstate reward expectation in RR (**Supplementary Figure 1C**), we hypothesized that this increased correlation between RR and UR2 could be related to the delayed development of reward expectation in RR. To test this, we correlated the activity of the first ten laps in RR with the last lap in R2 and the last lap in UR2. We found that the activity in the first few laps in RR were more correlated with the last lap of UR2 than with the last lap of R2 (average second lap correlation in RR when correlated with R2: 0.34 vs when correlated with UR2: 0.43, t = 5.638, p = 0.005, two-sided paired *t* test, **Supplementary Figure 3E**). However, with more laps in RR, the correlation with R2 increased. Nevertheless, after 10 laps in RR, this correlation was not significantly different than the correlation with UR2 (Lap 10 in RR, correlated with R2: 0.56 vs UR2: 0.57, t = 0.081, p = 0.939, two-sided paired *t* test, **Supplementary Figure 3E**). This suggests that after reward reinstatement, when reward expectation is still low, the spatial representation remains similar to when there was no reward and reward expectation was low. As reward expectation increases in RR, the representation starts to resemble the one when reward expectation was high, but it is not fully restored. Instead, the representation in RR is a combination of the most recent representation with low reward expectation and the more remote representation with high reward expectation.

### At the ensemble level, place cells form lower correlated networks across days when reward expectation is low

To better understand the relationships between reward expectation and across-day stability of place cells, we created weighted network graphs (Barabási et al., 2023; Bassett & Sporns, 2017; Humphries, 2018) based on place cell activity. In these graphs, nodes are place cells, and their across-day correlations are edges or links between the nodes. Specifically, we divided place cells defined in R1 or UR1, and constructed adjacency matrices for each graph by correlating the pairwise activity of these cells in R1 or UR1 with their activity on the following day in R2 or UR2, respectively. Therefore, the adjacency matrices for each of the graphs are matrices of pairwise correlation coefficients across days with dimensions equal to the number of nodes (place cells). This resulted in the generation of weighted network graphs, where each node represents a place cell defined in either R1 or UR1, and the edge linking any two nodes represents the Pearson correlation coefficient between the activity of all those cells with their activity in R2 or UR2 (see Methods).

To visualize the graphs, we utilized a force directed topographical layout (ForceAtlast2, Gephi, (Jacomy et al., 2014)). In this layout, nodes repulse each other like charged particles, while the edges attract the nodes like a spring. In a scenario where all place cells are stable, this topology will be spherical due to the circular nature of our track. The edge weights will attract place cells (nodes) with fields that are closer in location on the track together as they will have greater correlation and the nodes will push each other away from the center. We first correlated the activity of place cells in the first half of R1 with their activity in the second half of R1 (control-graph), where place cells are generally stable (Krishnan et al., 2022). Indeed, we found that the topology of the graph was circular, and the place cells or nodes with fields closer to each other on the track were arranged in a circular fashion, with very few crisscrossing edges (**Supplementary Figure 4A**). Such crisscrossing edges would indicate potential drift of a place cell since it is now correlated to another node with a different field location. Consequently, a dataset without any inherent correlation such as between R1 and shuffled timeseries will look collapsed without any apparent arrangement of the place cells or nodes by their location on the track (**Supplementary Figure 4B)**.

We visualized the weighted network graphs created between R1 and R2 (R-networks) as well as UR1 and UR2 (UR-networks). We observed that R-networks formed a spherical topology similar to our control graph (**Figure 3A-B**). Although the number of crisscrossing edges was higher than the control graph, indicating more drift across days than within a session, the nodes were arranged in a circular fashion based on their location along the track. We found that nodes in UR-networks were also arranged in a circular fashion but exhibited more crisscrossing than R-networks.

**Figure 3.**
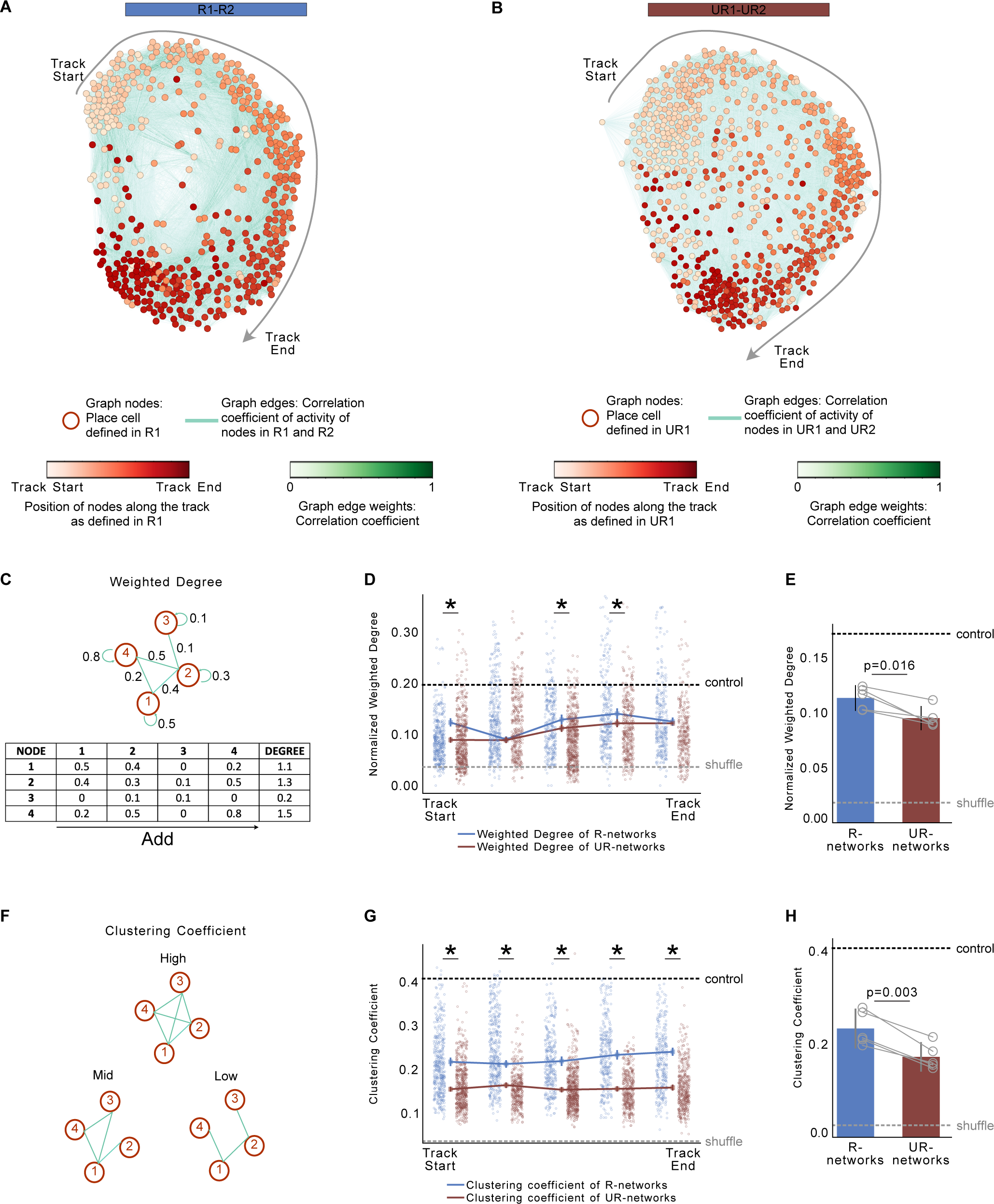
Place cells display stronger network correlation and connectivity across days when reward expectation is high. **(A)** Example network graphs of neuronal co-activity. All graphs are from the same mouse. Each node (in shades of red) represents a place cell defined in R1. Edge weights (in shades of green) represent Pearson correlation coefficient between the activity of nodes in R1 with their activity in R2 (R-networks). The graph is visualized using a Force Atlas 2 topology which arranges the nodes in a circular fashion grouping track positions that are closer to each other (see Methods). **(B)** Same as A but between UR1 and UR2 (UR-networks). **(C)** Depiction of weighted degree calculation which is measured as the sum of all edge weights of a node. (**D**) Nodes were divided by their location on the track (5 bins of 40 cm each). Normalized weighted degree was obtained by dividing the average weighted degree at each location bin by the total number of nodes in that bin. (**E**) Lines represent average weighted degrees from R-networks (blue) and UR-networks (red). Each dot is a node with place field in that track location in R1 (blue) or UR1 (red) (**F**) Average weighted degree plotted for each animal (n = 5 mice). (**F**) Depiction of clustering coefficients. A network with high clustering coefficient (=1) is one in which every node in the network is connected to each other by an edge. On the other hand, a network with low clustering coefficient only has fewer edges connecting the nodes. (**G**) Lines represent average clustering coefficient of nodes binned by track position. Each dot is a node with place field in that track location in R1 (blue) or UR1 (red). (**H**) Average clustering coefficient for each animal (n = 5 mice, grey circles). All error bars represent 95% confidence intervals. P-values were calculated using a Kolmogrov Smirnov (KS) test in D, G and a two-sided paired *t* test in E, H. In D, E, G, H, dotted lines represent average weighted degree and clustering coefficient obtained from control and shuffled networks, shown in **Supplementary Figure 4**.

To quantify this observation, we used two empirical measures. First, we used the weighted degree (**Figure 3C-E**), which measures the sum of all edge weights of a node (**Figure 3C**). This provides an estimate of aggregate network correlation. A higher weighted degree signifies greater ensemble correlation. Second, we used the clustering coefficient (**Figure 3F-H**), which measures the ratio of the number of edges among the nodes to the maximum number of edges possible in the node’s neighborhood (**Figure 3F**). This provides an estimate of local network connectivity. A higher clustering coefficient signifies greater stability within local networks among place cells with nearby field locations, while a low number signifies greater drift. We then binned the place cells or nodes by their location along the track (into 5 bins of 40cm each) and examined how the network varied along different regions of the track. As the number of place cells representing each location on the track varies in number (**Supplementary Figure 5**) and since the weighted degree scales with the number of nodes, we normalized the weighted degree by the number of nodes at each location.

We found that the normalized weighted degree and clustering coefficients were higher in the within-session network (control) than in the across-day networks (both R and UR-networks). This implies that ensembles are much more correlated within a session, and that there is considerable drift in the network across days, even when reward expectation is high (**Figure 3**, **Supplementary Figure 4**). However, we found R-networks to have higher weighted degrees and clustering coefficients than UR-networks (**Figure 3C-H**). Thus, R-networks have stronger network connectivity both locally (**Figure 3G-H**) and on aggregate (**Figure 3D-E**), than UR-networks. This indicates that place cells are more co-active across days as a network when reward expectation is high, and higher reward expectation better preserves network topology across days compared to low reward expectation. These findings confirm that high reward expectation can lead to greater stability at both the single cell and network levels.

We found these network parameters to generally be higher throughout the track in R-networks than UR-networks. However, the normalized weighted degree did not show statistical significance across all track locations. Surprisingly, despite the reduced number of cells near the rewarded location in UR1 (**Supplementary Figure 5**) and the absence of reward, we found that the weighted degrees of UR-networks were similar to R-networks around the rewarded location. We found, however, that R-networks had higher weighted degree in the beginning, as we observed at the single cell level (**Figure 2G**). Since the beginning and end of the track in our paradigm are contiguous, we posit that the reduction at the beginning may be a spillover effect from the lack of reward at the end.

### Place cells with high lap-by-lap reliability are more stable across days and are reduced in number when reward expectation is lowered

Next, to examine the relationship between place cells dynamics within a session and across-day stability, we assessed the lap-by-lap reliability of all place cells (Krishnan et al., 2022). Here, reliability combines two features of a place field: one, how often the cell fires in its place field across laps and two, how spatially precise the firing is on each lap with respect to the mean place field. We previously found that the lap-by-lap reliability of place cells decreased when reward expectation was lowered (Krishnan et al., 2022). Here, we first looked to see whether reliability remains low when reward expectation is lowered across days. Indeed, we found that place cell reliability was significantly lower in both UR1 and UR2 compared to R1, R2 and RR (**Supplementary Figure 6**). This was true across the population when place cells were defined in each of the conditions separately (**Supplementary Figure 6A-B**). Additionally, we were able to track place cells in R1 that had fields (not necessary with fields at the same position) across all the conditions (R2, UR1, UR2, RR), allowing us to see how the reliability of fields changes across conditions in the same cells. These common place cells revealed high reliability in R1 and R2 that decreased in UR1 and UR2 and increased in RR (**Supplementary Figure 6C**), similar to what is observed across the entire population. We next defined place fields with a reliability of >=0.5 as having high reliability, and the rest as having low reliability. We found that only 33 out of the 658 common place cells (5%) maintained place fields with high reliability throughout all conditions. Of these cells, the majority (30 cells) came from two mice (**Supplementary Figure 6D-F**). This shows that place field reliability is not a feature determined by cellular properties but is instead a flexible property that depends on the internal state of reward expectation.

We quantified the percentage of place cells with highly reliable fields and, unsurprisingly, found it was lower in both UR1 and UR2 conditions compared to R1 and R2 (**Figure 4A**). We then asked whether the low proportion of place cells with highly reliable fields that did exist in UR1 were associated with stable fields across days. Indeed, we found that place cells with highly reliable fields in UR1 were more stable across days compared to place cells with less reliable fields (**Figure 4B-E**). Interestingly, these place cells with highly reliable fields in UR1 demonstrated a similar level of stability across days as the place cells with highly reliable fields in R1 (**Figure 4F**).

**Figure 4.**
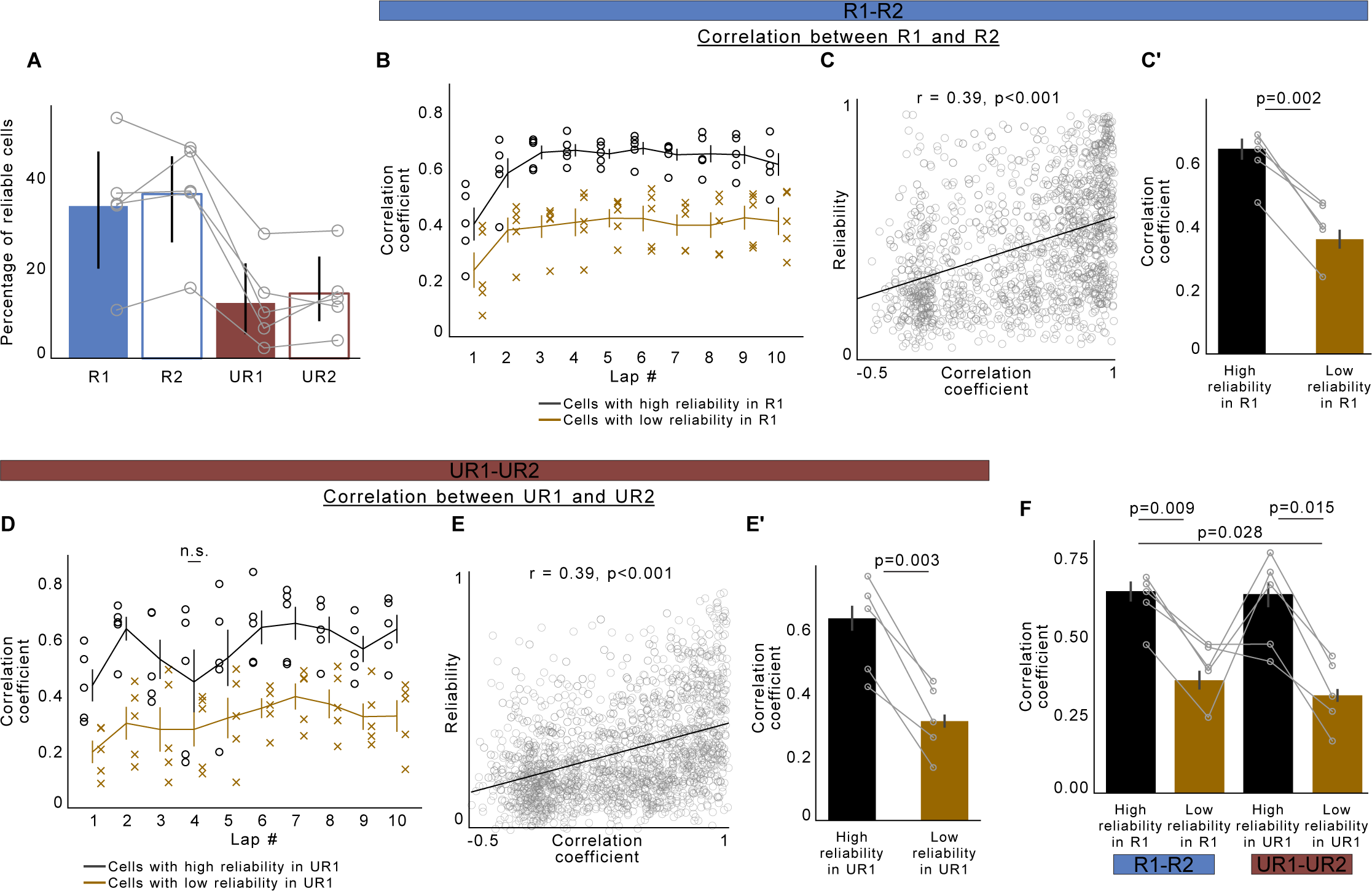
Place cells with more reliable fields from lap-to-lap are more stable across days. **(A)** Percentage of reliable cells across each condition. Dots represent individual mice. (**B**) Average correlation coefficient obtained from the activity of place cells in the last lap in R1 with their activity in the first 10 laps of R2. Circles indicate individual mice. Place cells were divided based on their reliability in R1. High reliable cells have reliability >=0.5 and low reliable cells have reliability < 0.5. **(C)** Scatter plot of the reliability of a place cell in R1 (y-axis) and the correlation coefficient between the mean place fields of the place cell in R1 and R2 (x-axis). Each circle is a place cell defined in R1. Black line shows linear regression fit. r and p-values were obtained using Pearson’s correlation coefficient. **(C’)** Animal-wise averages of the data in C divided based on the reliability of the place cell in R1. Circles indicate individual mice. P-values were obtained using a two-sided paired *t* test. **(D-E)** Same as B-C but between UR1 and UR2. (**F)** Same data as in C’ and E’, displayed together. P-values were obtained using a two-sided paired *t* test and corrected for multiple comparisons. In B and D, P-values (two-sided paired *t* test) were obtained between correlation coefficients of place cells with high reliable (black) vs low reliable (gold) fields. non-significant (n.s.) P-values are indicated, all other relationships were significant. All error bars indicate 95% confidence intervals. For full details on statistics and p-values, see **supplementary file 1**

Next, we examined the lap-by-lap correlation between the last lap of the previous day (in R1 and UR1) and the first ten laps of the next day (in R2 and UR2) for both the cells with high and low reliable fields, as previously described (**Figure 2H-I**). On the very first lap of R2 and UR2, place cells with highly reliable fields on the previous day, showed a greater correlation than cells with less reliable fields (correlation coefficient first lap R2: cells with high reliability: 0.40 ± 0.15, cells with low reliability: 0.24 ± 0.16. UR2: cells with high reliability: 0.44 ± 0.15, cells with low reliability: 0.20 ± 0.10). In place cells with highly reliable fields, this correlation also increased as mice ran more laps in both R2 and UR2 (**Figure 4B, D**, correlation coefficient first 2-10 laps R2: 0.64 ± 0.03, UR2: 0.59 ± 0.06). The correlation increased in cells with less reliable fields but to a much smaller extent and always remained lower than the correlation in cells with highly reliable fields (correlation coefficient first 2-10 laps R2: 0.41 ± 0.03, UR2: 0.33 ± 0.05, p < 0.05 two-sided paired *t* test). Among cells with highly reliable fields, the average correlation coefficient between R1 and R2 vs UR1 and UR2 was not statistically different (p > 0.05, two-sided paired *t* test, **Figure 4F**). We also found cells with highly reliable fields in R1 and UR1 to be more correlated at the network level compared to cells with less reliable fields (**Supplementary Figure 8**). These results confirm a recent finding that place cells with highly reliable fields are less prone to drift (Chiu et al., 2023). Additionally, cells with highly reliable fields in R1 were more stable in RR than cells with low reliable fields in R1. This is noteworthy considering RR was separated by two days from R1 and succeeded sessions where reward expectation was low (**Supplementary Figure 7D-E**). Thus, in an unchanging spatial environment, the proportion of highly reliable place fields determines the extent of representational drift across days. This feature is flexible and governed by the animal’s internal state.

## Discussion

In this study, our goal was to examine whether lowering reward expectation over consecutive days would increase drift across days in CA1. Our paradigm allowed us to record from the same group of hippocampal neurons over a span of three days as we changed the internal state of reward expectation of the animal by removing and reintroducing reward in the same spatial environment. Our findings revealed that both at the level of single cells and as a network, place cells exhibited greater drift of their fields when reward expectation was low across consecutive days, compared to when reward expectation was high. These results follow our previous study where we showed that reward expectation enhances the spatial encoding ability of hippocampal place maps (Krishnan et al., 2022). An important finding from our work is that cells with reliable place fields on a lap-to-lap basis within a session were more stable across days, regardless of whether reward expectation was high or low. The association between the reliability of place fields from one trial to the next and the stability of these fields over time has been reported in previous studies (Chiu et al., 2023; Sheffield & Dombeck, 2015). Our current results affirm this relationship and show that the increased drift when reward expectations are lowered is attributed to a decrease in the proportion of reliable place fields across the population of place cells.

The decrease in lap-to-lap reliability in CA1 due to lowered reward expectation is likely caused by a reduction in dopaminergic input from the Ventral Tegmental Area (VTA) (Krishnan et al., 2022). One of the known roles for hippocampal dopamine is in regulating synaptic transmission and dendritic excitability (Edelmann & Lessmann, 2018; Hansen & Manahan-Vaughan, 2014; Huang & Kandel, 1995; Lisman & Grace, 2005; Tritsch & Sabatini, 2012; Wiescholleck & Manahan-Vaughan, 2014), and high dendritic excitability is associated with higher reliability and long-term stability of place cells (Sheffield et al., 2017; Sheffield & Dombeck, 2015, 2018). Studies have also demonstrated that a decrease in dopamine leads to a reduction in across-day stability of place cells and increased drift (Kentros et al., 2004; Mamad et al., 2017; Martig & Mizumori, 2011; McNamara et al., 2014). Therefore, reduction of dopamine released in the hippocampus from the VTA with lowered reward expectation likely contributes to the observed reduction in lap-to-lap reliability and subsequent instability of place fields across days. Dopamine may accomplish this by enhancing the plasticity mechanisms that drive the reliability and stability of salient memories (Bittner et al., 2017; Madar et al., 2023; Plitt & Giocomo, 2021; Plitt et al., 2023; Thomas et al., 2023).

Dopamine release during periods of heightened reward expectation may indirectly enhance the stability of place maps by strengthening functional network correlations in CA1. Through network analysis, we found that place cells with proximate fields established correlated functional networks over consecutive days. This correlation was more prominent when the expectation of reward was higher, coinciding with elevated dopamine release in CA1. Given the limited glutamatergic reciprocal connectivity across CA1 pyramidal cells, heightened activity correlations among CA1 cell ensembles might be driven by reciprocal connections with interneurons (Geiller et al., 2023). Notably, CA1 interneurons express dopamine receptors (Puighermanal et al., 2017), suggesting a potential pathway for dopamine to modulate functional networks in CA1. Further investigation is needed to determine the specific impact of dopamine on CA1 network activity. Additionally, other neuromodulators, such as acetylcholine (Hasselmo, 2006; Mau et al., 2020), may also contribute to these processes and warrant further exploration.

Although lap-by-lap reliability decreased with lowered reward expectation, we observed that a small subset of place cells maintained highly reliable fields that remained stable across days. However, the proportion of place cells with high reliability was much smaller compared to when the reward expectation was high (Krishnan et al., 2022). Despite this, these cells were just as effective in maintaining their fields across days as the highly reliable cells that were present when reward expectation was high. The smaller proportion of highly reliable cells when reward expectation was low suggests that the CA1 may represent an impoverished memory of the environment compared to when reward expectation is high. Consequently, accurate retrieval of this memory may be limited by the reduced stability of these unreliable place fields.

Mechanistically, the presence of a small subset of reliable and stable place fields during low reward expectation could be attributed to residual levels of dopamine still present under this condition. The hippocampus receives dopamine from two sources: the VTA and the locus coeruleus (LC) (Kempadoo et al., 2016; Takeuchi et al., 2016). While VTA dopamine axons in CA1 exhibit decreased activity when reward expectation is lowered, inputs from the locus coeruleus to CA1 are unaffected by reward expectation (Heer & Sheffield, 2023) and have been shown to play a role in shaping contextual memories (Chowdhury et al., 2022; Kaufman et al., 2020). Therefore, it is plausible that the presence of a small number of reliable and stable place fields in the hippocampus during low reward expectation is mediated by inputs from the locus coeruleus. Additionally, other neuromodulators may also play a role in maintaining reliable place fields in the hippocampus (Hasselmo, 2006; Luchetti et al., 2020; Palacios-Filardo & Mellor, 2019). Future research should investigate these different neuromodulators and their influence on place field reliability as well as their activity with changing internal states. This can provide further insights into the mechanisms that enhance memory stability and reduce drift.

It is generally thought that once place fields are formed during an experience, they are strengthened through offline reactivation and retrieved upon re-exposure to the same environment (Carr et al., 2011; Ólafsdóttir et al., 2018; O’Neill et al., 2008; Skaggs & McNaughton, 1996; Wilson & McNaughton, 1994), a property that is reliant on dopaminergic input to the hippocampus from the VTA (McNamara et al., 2014). However, most studies focus on correlating the average activity of place cells across sessions. Here, we show that when checking the correlation on a lap-by-lap basis, the retrieval of previous spatial representations does not happen right away, even in a familiar environment without any changes to behavior or internal state (**Figure 2I, Supplementary Figure 3A**). We observed that the correlation between place cell activity across days was the lowest on the very first lap on day 2 in both the R and UR conditions. As the animal completed more laps, the correlation increased starting from the second lap and stabilized across subsequent laps. This suggests that the retrieval of the previous day’s place map does not happen immediately. Instead, it requires experience to reactivate the same input patterns and re-engage place cells to fire at the same location where they were previously active, maybe as the animal gradually remembers the context in which it is in. However, the lower correlation in the very first lap could also be attributed to other changes in the animal’s internal state. It is possible that animals display more caution when placed into a virtual environment, even with extensive training. Although we found no differences in lap velocity across days (**Supplementary Figure 3A**), we did not measure other parameters, such as the pupil diameter, which could indicate differences in arousal (Bradley et al., 2008) and potentially affect place cell firing (Bradley et al., 2008; Krishnan et al., 2022; Pettit, Yuan, et al., 2022). Further investigation is needed to determine if there are other behavioral differences or variations in internal states during the initial laps compared to the later laps that could contribute to this delay in representation reinstatement.

It is worth noting that the average correlation on the first lap was lower (0.39) than other laps (0.61), but not zero. This was also true when considering just the place cells with highly reliable fields on the previous day (first lap: 0.42, other laps: 0.62). The place cells that did reactivate on the very first lap could be the ones that were preferentially strengthened over others. These place cells could help recruit the other cells into the map as the animal experiences the environment. CA1 is thought to integrate the animal’s current experience, represented by inputs from the entorhinal cortex, with representations of past experiences supplied by the direct inputs from CA3 or indirectly from the Dentate Gyrus (DG) (Brun et al., 2002; Grienberger & Magee, 2022; Keinath et al., 2020; Mizumori et al., 1999; Plitt & Giocomo, 2021). It could be that the very first lap is dominated by inputs potentiated by the entorhinal cortex, followed by the CA3 and DG as the animals experience the context and match the current experience with the past. This hypothesis is further supported by evidence that reliable cells in both R and UR conditions on day 1 demonstrated similar retrieval dynamics with experience on day 2. However, it is important to highlight that the average correlation across days, even among place cells with highly reliable fields was approximately 0.6, indicating that the representation doesn’t fully return to the previous representation and contains inherent drifts despite similar internal states, as shown in previous studies in CA1 (Hainmueller & Bartos, 2018; Kinsky et al., 2018; Lee et al., 2020; Rubin et al., 2015; Ziv et al., 2013). This also implies that it is important to consider lap-by-lap dynamics when quantifying the amount of drift across days and averaging the activity of place cells across the session may skew the amount of drift versus stability quantified within a session.

Finally, we found that prior representations were not fully restored following reward-reinstatement. This supports our previous finding where we found reward-reinstatement on the same day following reward expectation extinction also did not fully restore prior representations (Krishnan et al., 2022). Interestingly, we found that the place map following reward-reinstatement was correlated with both the recent representation in UR2 with a different internal state and the remote representation in R2 with the same internal state. This suggests that while CA1 chunks external events based on different internal states, the cells that represent each chunk of the experience combine, potentially creating a representation that holds in memory the different events that occurred in that environment (Smith & Mizumori, 2006). However, whether this combined representation continues to remain stable on subsequent days needs to be determined. Since reward reinstatement increases the number of reliable place cells (**Supplementary Figure 6A**) and restores dopaminergic input to the hippocampus (Krishnan et al., 2022) it is likely that a more stable representation will emerge over multiple days following reward reinstatement.

In conclusion, our findings demonstrate that in an unchanging spatial environment the extent of representational drift in the CA1 region across time is in-part determined by the animal’s internal state of reward expectation. When reward expectation is high, VTA dopamine levels in CA1 are also high and this increases the proportion of reliable place fields which are more likely to be stable across time, thus reducing representational drift. This tight interconnection between place field reliability and stability and the brain’s ability to modulate the proportion of these cells through neuromodulatory circuits provides a mechanism to control the extent of drift in the hippocampus, thus controlling the strength of memory encoding and retrieval.

## Acknowledgements

We thank Chery Cherian for his help with initial surgeries and data collection. We thank Matt Rosen for his assistance with the network analysis. We thank Douglas GoodSmith, Chad Heer and Matt Rosen for feedback on the manuscript.

## Author contributions

S.K. and M.S. conceived and designed the experiments. S.K. collected and analyzed the data. S.K. and M.S. interpreted the data and wrote the manuscript. M.S. supervised the research and obtained funding.

## Data availability

Raw imaging data is large and not feasible for upload to an online repository but is available upon request to the lead contact at. Processed source data for all figures and associated statistical analysis are provided with the paper.

## Code availability

Scripts used for data analysis are available on Github (https://github.com/seethakris/RewardMultiDay)

## Methods

### Subjects

All experimental and surgical procedures were in accordance with the University of Chicago Animal Care and Use Committee guidelines. The mice were individually housed in a reverse 12-hour light/dark cycle, and the behavioral experiments were carried out during the animal’s dark cycle. For this study, we used 10–12-week-old male C57BL/6J wildtype (WT) mice. Male mice were used over female mice due to the size and weight of the headplates (9.1 mm x 31.7 mm, ∼2g) which were difficult to firmly attach on smaller female skulls. At this age, females were generally smaller than males and their weight was further reduced by water restriction (∼15g in females compared to ∼22g in males). Due to their smaller weight following water-restriction, females had difficulty being head-restrained, running on the treadmill, and learning the task at the required speed before viral over-expression.

### Mouse surgery and viral injections

Mice were anaesthetized (∼1%-2% isoflurane) and injected with 0.5 ml of saline (intraperitoneal injection) and 0.5 ml of Meloxicam (1-2 mg/kg, subcutaneous injection) before being weighed and mounted onto a stereotaxic surgical station (David Kopf Instruments). A small craniotomy (1-1.5 mm diameter) was made over the hippocampus (1.7 mm lateral, -2.3 mm caudal of Bregma). For population imaging of hippocampal CA1 cells, a genetically-encoded calcium indicator, AAV1-CamKII-GCaMP6f (pENN.AAV.CamKII.GCaMP6f.WPRE.SV40 was a gift from James M. Wilson – Addgene viral prep #100834-AAV1; http;//n2t.net/addgene:100834 ; RRID:Addgene_100834) was injected (∼50 nL at a depth of 1.25 mm below the surface of the dura) using a beveled glass micropipette leading to GCaMP6f expression in a large population of CA1 pyramidal cells. Mice were separated into individual cages and were water restricted from the following day (0.8-1.0 ml per day). After a week of water restriction, mice underwent another surgery to implant a hippocampal window (Dombeck et al., 2010). Following implantation, the head-plate was reattached with the addition of a head-ring cemented on top of the head-plate which was used to house the microscope objective and block out ambient light. Post-surgery mice were given 2-3 ml of water/day for 3 days to enhance recovery before returning to the reduced water schedule (0.8-1.0 ml/day). Expression of GCaMP6f reached a somewhat steady state ∼20 days after the virus was injected.

### Behavior and Virtual Reality

The virtual reality (VR) environments that the mice navigated through were created using VIRMEN (Aronov & Tank, 2014) and were rich in visual cues. Mice were head restrained with their limbs comfortably resting on a freely rotating Styrofoam wheel (‘treadmill’). Movement of the wheel caused movement in VR via a rotatory encoder that detected treadmill rotations and fed this information into a computer. Mice received a water reward (4 µl) through a waterspout upon completing each traversal of the 2m long linear track (a lap), which was associated with a clicking sound from the solenoid. Licking was monitored by a capacitive sensor attached to the waterspout. Upon receiving the water reward, a short VR pause of 1.5 s was implemented to allow for water consumption. Mice were then virtually teleported back to the beginning of the track and could begin a new lap traversal. Mouse behavior (running velocity, track position, reward delivery, and licking) were collected using a PicoScope Oscilloscope (PICO4824, Pico Technology, v6.13.2). Behavioral training to navigate the virtual environment began 4-7 days after imaging window implantation (∼30 minutes per day) and continued until mice reached >4 laps per minute, which took 10-14 days. Although, some mice never reached this level. This high level of training was necessary to ensure mice continued to traverse the track similarly after reward was removed from the environment (Krishnan et al., 2022). Initial experiments showed that mice that failed to reach this criterion typically did not traverse the track as consistently without reward across days. Such mice were not used for imaging. Additionally, since we are testing changes in reward expectation, only animals that displayed pre-licking in the familiar environment before reward delivery were used for imaging. The rate of success in training mice to reach this criterion was ∼60%. In mice that reached criteria, imaging commenced the following day.

### Two-photon imaging

Imaging was done using a laser scanning two-photon microscope (Neurolabware). Using an 8kHz resonant scanner, images were collected at a frame rate of 30 Hz with bidirectional scanning through a 16x/0.8 NA/3 mm WD water immersion objective (MRP07220, Nikon). GCaMP6f was excited at 920 nm with a femtosecond-pulsed two photon laser (Insight DS+Dual, Spectra-Physics) and emitted fluorescence was collected using a GaAsP PMT (H11706, Hamamatsu). The average power of the laser measured at the objective ranged between 50-70 mW. A single imaging field of view (FOV) between 400-700 µm equally in the *x/y* direction was positioned to collect data from as many CA1 pyramidal cells or dopaminergic axons as possible. Time-series images were collected through Scanbox (v4.1, Neurolabware) and the PicoScope Oscilloscope (PICO4824, Pico Technology, v6.13.2) was used to synchronize frame acquisition timing with behavior.

### Imaging Sessions

All experimental sessions were multi day imaging sessions, and the same cells were imaged across all days. The environment used during the imaging sessions was the same environment that the animals trained in. Each imaging session lasted ∼8-12 minutes and was always presented in the same order. On Day 1, mice (n = 5) were exposed only to the familiar rewarded environment (R1). At the end of the first imaging session, a 1-minute time-series movie was collected at a higher magnification and then averaged to aid as a reference frame in finding the same imaging plane on subsequent days. On these days, it took about 10 minutes on average to locate the same imaging plane as before during which time mice were shown a dark VR screen. On Day 2, mice first experienced the rewarded environment (R2) followed by the unrewarded environment (UR1). On Day 3, unrewarded environment (UR2) was experienced first followed by the rewarded environment again (Re-Rewarded, RR).

### Image Processing and ROI selection

Time-series images were preprocessed using Suite2p (Pachitariu et al., 2017). Movement artifacts were removed using rigid and non-rigid transformations and assessed to ensure absence of drifts in the *z*-direction. Datasets with visible *z*-drifts were discarded (n = 2). Imaging planes acquired from each day were first motion corrected separately. ImageJ (NIH) was then used to align the motion corrected images relative to each other by correcting for any rotational displacements. The images across all days were then stitched together and motion corrected again as a single movie. Regions of interest (ROIs) were also defined using Suite2p and manually inspected for accuracy. Baseline corrected ΔF/F traces across time were then generated for each ROI and filtered for significant calcium transients (Sheffield et al., 2017). Rastermap was used to display the correlated patterns of fluorescence in cells across all conditions (Stringer et al., 2023).

### Identifying when pre-licking stops

Licking data was collected using a capacitive sensor on the waterspout. As previously described (Krishnan et al., 2022), mice displayed anticipatory licking behavior in the rewarded environment, which continued in UR1 before decaying exponentially. The lap where pre-licking stops in UR1 was defined as the lap following 2 consecutive laps with an absence of these anticipatory licks (Krishnan et al., 2022). None of the mice displayed pre-licking in UR2. Only laps that followed the end of anticipatory licking were used for analysis in UR1.

### Position decoding

A naïve Bayes decoder (scikit-learn, v1.0, (Pedregosa et al., 2011)) was used to decode the animal’s position on the track from the activity of all cells. The decoder was trained and tested exactly as described previously (Krishnan et al., 2022). The training data consisted of 60% of laps (randomized) in R2 in each animal and the decoder was tested on the remaining laps in R2 as well as on laps in UR1 before and after pre-licking stopped. Quality of decoder fit was assessed by calculating the coefficient of determination (R^2^) between the actual location of the animal and the location predicted by the decoder.

### Place fields and place field parameters

Place fields were defined as previously described (Krishnan et al., 2022). Briefly, the 2 m track was divided into 40 position bins (each 5 cm wide). The running behavior of the animal was filtered to exclude time periods where the animal was immobile (speed <1 cm/s). Filtering was done to ensure that place cells were defined only during active exploration. Extracted place fields satisfied all the following criteria 1) were >10 cm in width. 2) with average ΔF/F>10% above baseline 3) had average ΔF/F within the field was >4 times the average outside the field 4) The cell displayed calcium transients on >30% of the laps 5) P-value from shuffling the data 1000 times was <0.05. The ΔF/F of each place field was binned into laps by space (40 bins) matrix. The mean place field was calculated by averaging the ΔF/F across laps. The following parameters were extracted from each place field:

### Lap-by-lap Reliability

To calculate reliability, we computed the Pearson correlation between each lap traversal to obtain a lap × lap matrix. To obtain the reliability index, the average of this correlation matrix was multiplied by the ratio of number of laps with a significant calcium transient within the field and the total number of laps. The reliability index is 1.0 if the cell fires at the same location in each lap and 0.5 if it fires at the same position but only in half the laps, and so on. We used the threshold of 0.5 to classify cells as being high (>=0.5) or low (<0.5) in reliability. We also tested 0.6 as a threshold for defining highly reliable place fields without observing much change in the results (**Supplementary Figure 7B-C**). When we used higher thresholds (>=0.7), we found that some animals had only a few or no place cells with highly reliable fields in UR1, which made accurate statistical comparisons difficult. As a result, we determined that a threshold of 0.5 was the most suitable for distinguishing place cells with high and low reliable fields.

### Center of Mass (COM)

The COM for a place field was calculated as,

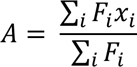

Where F is the ΔF/F in each bin *i* and xi is the distance of bin *i* from the start of the track. The mean COM was calculated by averaging across all laps.

### Population vector correlation

To determine the level of similarity in representations in different conditions, we calculated a mean Population Vector (PV) correlation. For each spatial bin, for all cells (not just place cells), we defined the PV as the average ΔF/F for that neuron in that bin across all lap traversals. We calculated the PV correlation in one condition with that of the matching position in another condition and averaged correlations across all positions (Figure 2 E-G).

### Lap-wise spatial correlation

Place cells were defined in one condition and their activity in the last lap was correlated (using Pearson Correlation) with the activity in the first ten laps on the next day.

### Network Analysis

Network analysis was performed on extracted place cells. Place cells (N) were defined in R1 and UR1. Pairwise Pearson’s correlation coefficient was calculated between mean place field activity in R1 and UR1 and mean place field activity on R2 and UR2 to form a NxN adjacency matrix. We also calculated an adjacency matrix from correlation coefficients between two halves of the session in R1 (Control), and between R1 and shuffled place cell activity in R2 (shuffle). Correlation coefficients that were negative and not statistically significant were removed. Note that the place cells are being correlated to their activity on the next day and will have an autocorrelation of 1 only if their place fields reappear on the same location the next day. Place cells were nodes and weights of the edges between two nodes were the correlation coefficients. Networks were plotted using the open-source tool Gephi (https://gephi.org/) and network topology was visualized using Force Atlas2. Network parameters (weighted degree and cluster coefficients) were calculated using the networkx package in Python (3.8.5). To display the weighted degree and cluster coefficients of the nodes across track position, we binned nodes by their center of mass on the track, each bin being 25 cm in length.

### Statistics

For unrelated samples from different groups, we performed a Kolmogrov Smirnov (KS) test. For related samples, we performed a paired t-test. Multiple comparisons of related samples were corrected with Bonferroni post-hoc. Detailed statistics including n, test-statistic and p-values for each figure are provided in Supplementary file 1 and where possible actual p-values and data distribution have been displayed. Boxplots are plotted to display the full distribution of the data. The box in the boxplot ranges from the first quartile (25^th^ percentile) to the third quartile (75^th^ percentile) and the box shows the interquartile range (IQR). The line across the box represents the median (50^th^ percentile). The whiskers extend to 1.5*IQR on either sides of the box and anything above this range is defined as an outlier. Additionally, estimation statistics were used to ascertain the level of differences between distributions by using the DABEST (v0.3.1, Data Analysis with Bootstrap-coupled Estimation) package (Ho et al., 2019). Estimation plots display the median difference between two conditions against zero difference, with error bars displaying 95% confidence intervals of a bootstrap generated difference (5000 resamples). A kernel density fit (shaded curve) on the resampled difference is also displayed alongside. This difference was compared against zero. Correlation coefficients were obtained using Pearson’s correlation coefficient. Data preprocessing and analysis was done on MATLAB (Mathworks, Version R2022a) and Python 3.8.5 (https://www.python.org/).

