## Supplementary Figure for "Reward Expectation Reduces Representational Drift in the Hippocampus"

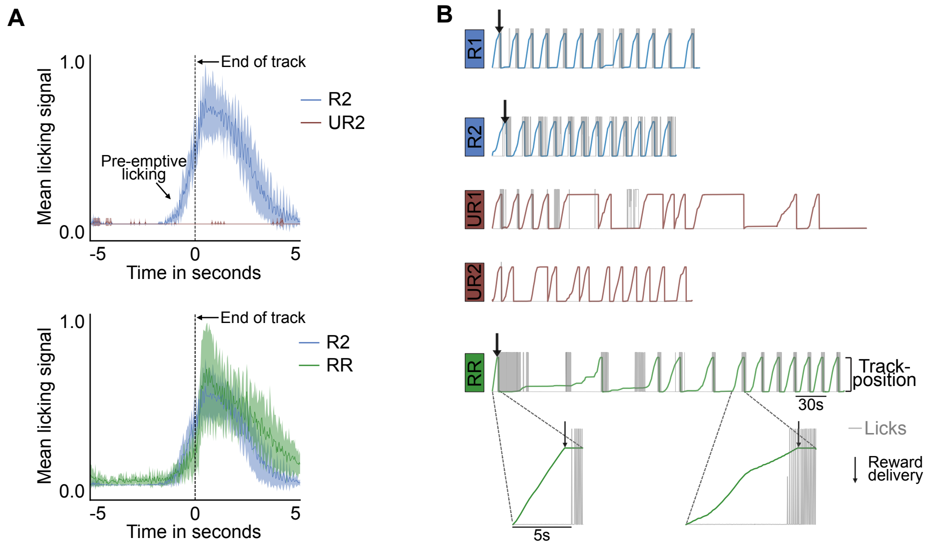


**Supplementary Figure 1: Licking behavior with reward expectation** (A) Mean number of licks around reward delivery (time=0). Unrewarded (UR2) and re-rewarded (RR) conditions are compared with the rewarded condition on day 2 (R2). Animals (n=5 mice) displayed pre-emptive licking in R2 (blue) and RR (green) but not in UR2 (red). Number of licks were calculated for each mouse on each lap and were binarized as 1 or 0 depending on if the animal licked or not in the given time bin. Shading represents standard error of the mean (s.e.m). (B) Example behavior traces from one mouse across all conditions. The first 12 laps are shown for each condition. Licking behavior is shown in grey. Water reward is delivered at the end of the track in R1, R2 and RR conditions, indicated by a black arrow only in the first lap for simplicity but a reward was given on each lap in these conditions. No water reward was given in UR1 and UR2. Reward expectation remained diminished in UR2 and this animal did not pre-emptively lick in UR2. The zoomed in laps in RR show that the animal licked after reward delivery in lap 1 when reward was reinstated but showed pre-emptive licking by lap 6.

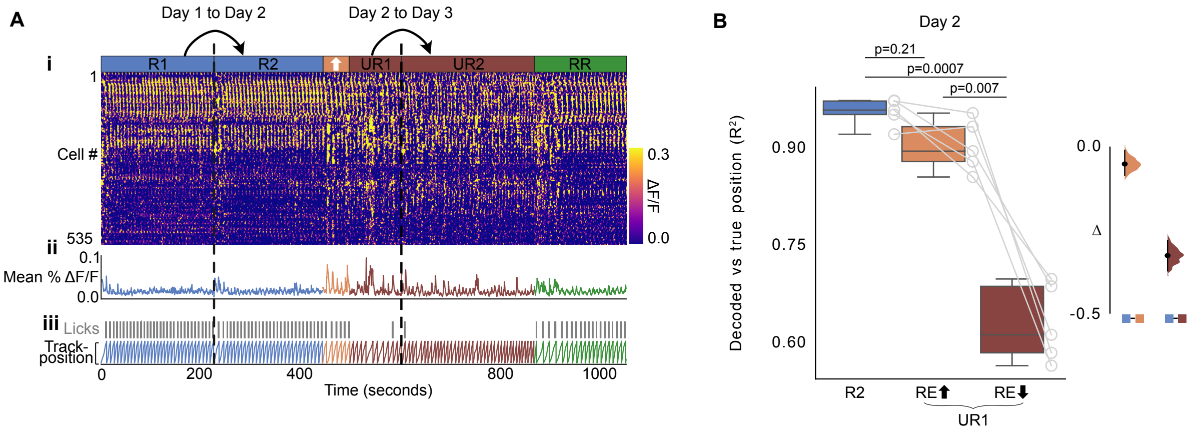


**Supplementary Figure 2: Neural activity with reward expectation.** (A) i: Rasterplot (Stringer et al., 2023) representing fluorescence changes (Δ*F*/*F*) of cells in A across time. Cells along the y-axis are arranged with the most correlated cells next to each other. ii: Mean Δ*F*/*F* of the cells in (i). iii: Mouse licking behavior. iv: Mouse track position. As defined previously (Krishnan et al., 2022), laps before animal stops consistently licking in UR1 were considered laps with high reward expectation ($RE_{high}$, orange laps) and after licking stops are laps with low reward expectation ($RE_{low}$, brown laps, see Methods). (B) (Left) Boxplots (see methods for definition) show distribution of mean decoder $R^2$ in the different conditions on Experimental Day 2. $RE_{high}$ and $RE_{low}$ represent laps in UR1 with high and low reward expectation respectively. These laps were defined based on when each animal stopped pre-licking (see methods). Decoder was trained on initial laps in R2 and tested on remaining laps in R2 and UR1 (see Methods). Circles represent individual animals. *P* values were obtained using a two-sided Paired *t* test with Bonferroni Correction for multiple comparisons (Right) Bootstrapped mean differences (Δ) with 95% Confidence Intervals (CI) (error bar). *X*-axis indicates the comparisons made.

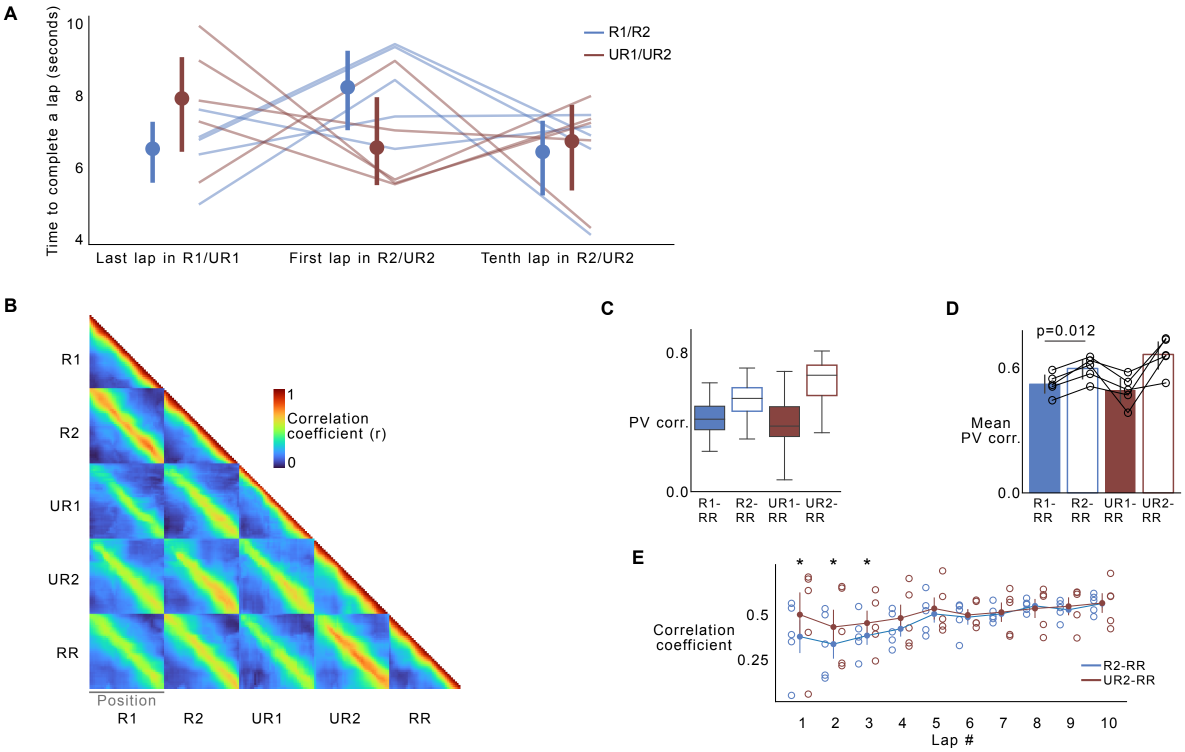


**Supplementary Figure 3. Reward reinstatement does not fully restore the original representation when reward expectation was high.** (A) Time taken to complete the last lap (left) in R1 (blue) and UR1 (red) compared to the first lap (center) and tenth lap (right) in R2 (blue) and UR2 (red). None of the relationships were significantly different (two-sided paired *t-test*) (B) Position vector (PV) correlation between all pairs of position across all conditions. The correlation was averaged across all imaged cells (n=6880 cells) and all animals (n=5 animals). (C) Boxplot showing distribution of PV correlations across all cells and all conditions with RR. All relationships were statistically significant. For full details on statistics and p-values, see supplementary file 1. (D) Animal-wise averages of data in C. Circles display individual animals. Significant P-value obtained using two-sided paired *t* test is displayed. (E) Average correlation coefficient across all place cells defined in R2/UR2 and their correlation with the first 10 laps in R2 (blue) and UR2 (red) respectively. Circles indicate individual mice. Asterix (*) denote significant *P* values (two-sided paired *t* test, $P < 0.01$) obtained by comparing R2-RR correlation (blue) with UR2-RR correlation (red) at each lap. Error bars in D and E indicate 95% confidence intervals. For full details on statistics and p-values, see **supplementary file 1**.

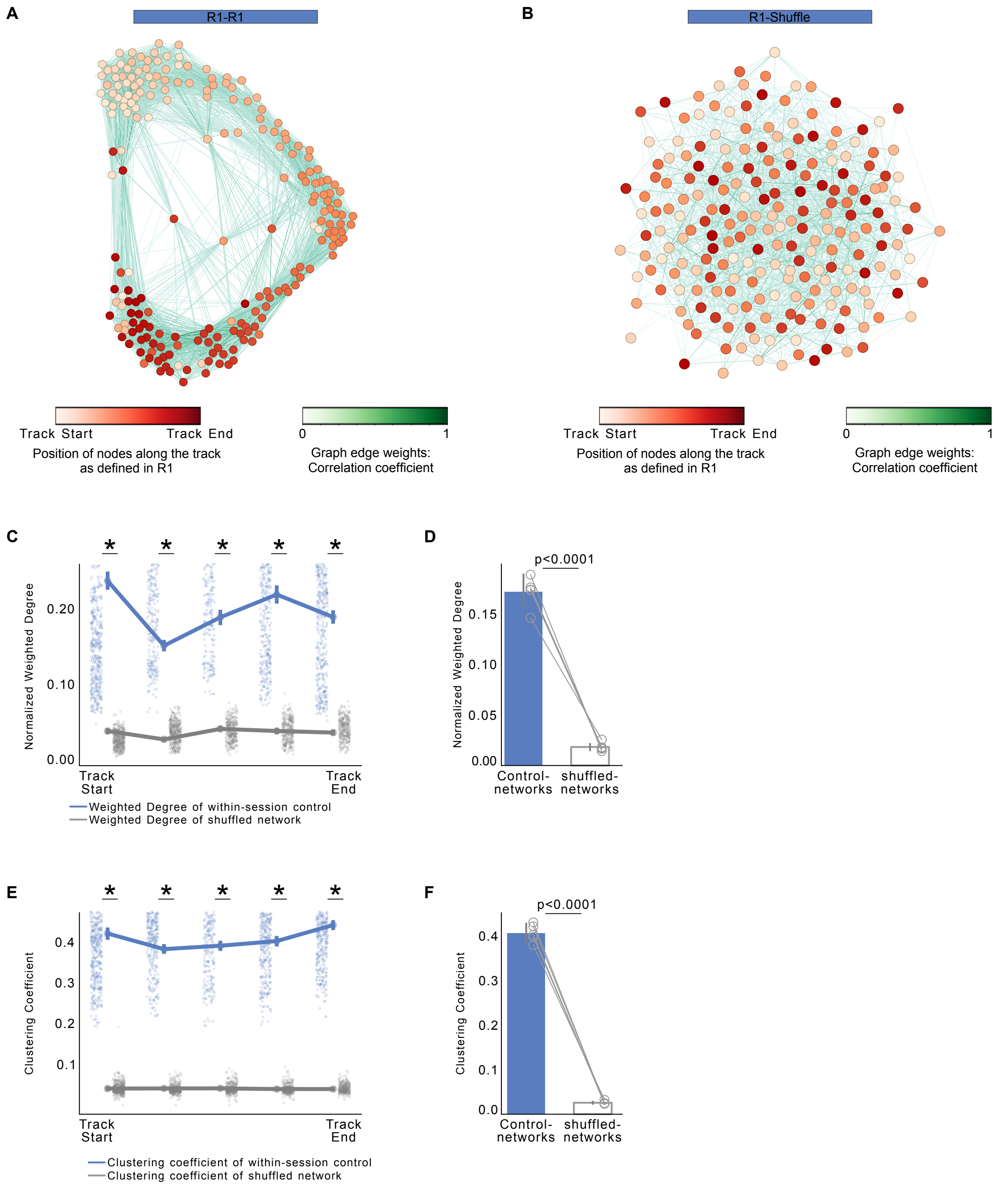


**Supplementary Figure 4. Network graphs showing within-session networks and shuffled networks. (A-B)** Example network graphs of neuronal co-activity. Graphs are from the same mouse as **Figure 3**. Each node (in shades of red) represents a place cell defined in R1. Both graphs are visualized using a Force Atlas 2 topology. (**A**) Edge weights (in shades of green) represent Pearson correlation coefficient between the activity of nodes in the first half of R1 with their activity in the second half of R1 (control-network). (**B**) Edge weights were calculated between activity of nodes in R1 with a shuffled timeseries (shuffled-network). (**C**) Nodes were divided by their location on the track (5 bins of 40 cm each). Normalized weighted degree was obtained by dividing the average weighted degree at each location bin by the total number of nodes in that bin. Lines represent average weighted degrees from control-networks (blue) and shuffled-networks (grey). Each dot is a node with place field in that track location in R1 (**D**) Average weighted degree plotted for each animal (n = 5 mice). (**E**) Lines represent average clustering coefficient of nodes binned by track position. Each dot is a node with place field in that track location in R1. (**F**) Average clustering coefficient for each animal (n = 5 mice). All error bars represent 95% confidence intervals. P-values were calculated using a Kolmogrov Smirnov (KS) test in C, E and a two-sided paired *t* test in D, F.

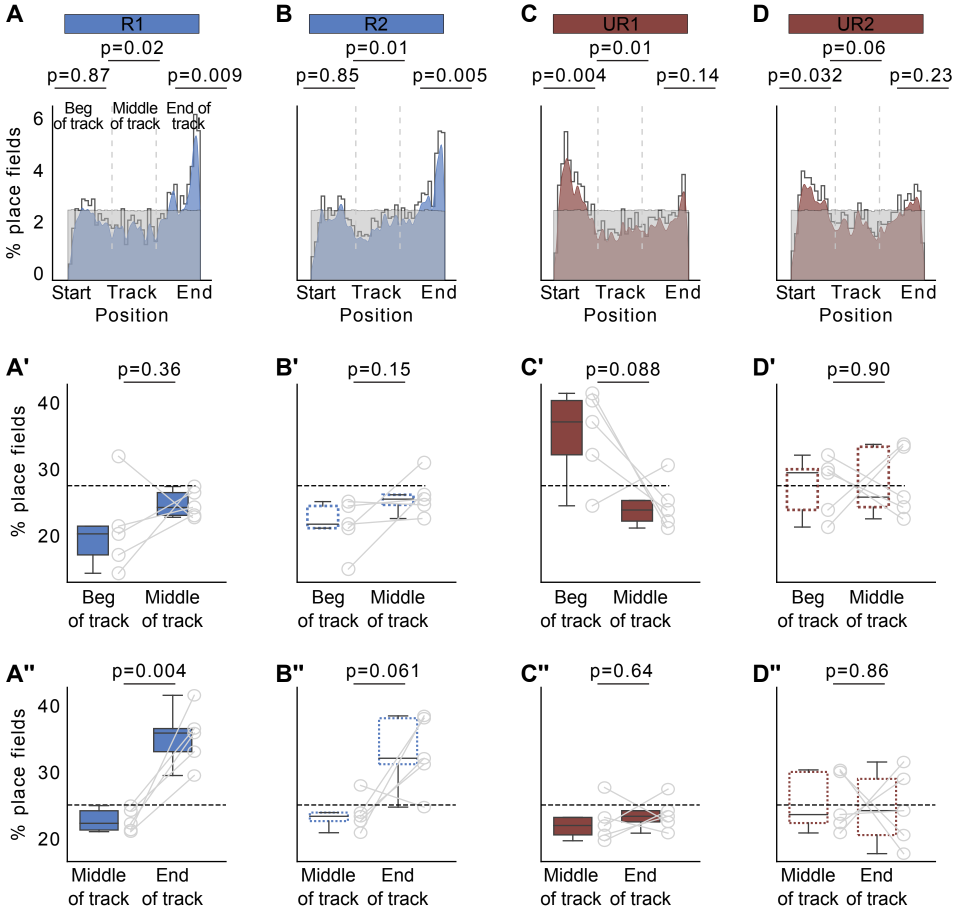


**Supplementary Figure 5. Distribution of place fields by location across each condition. (A-D)** Distribution of place field center of mass (COM) locations in each condition (displayed on top) pooled from all mice ($n = 5$ mice). Plots show observed density (gray line), uniform distribution (gray shade) and Gaussian distribution of place field density (color). *P* values (two-sided *t* test) were obtained by calculating the place field distribution within each zone (Beginning, middle, end – indicated by dashed vertical lines) with the uniform distribution. (**A'-D'**) Percentage of place fields in the beginning (Beg) of the track versus middle of the track in each animal (circles). (**A''-D''**) Percentage of place fields in the middle of the track versus end of the track in each animal (circles). Horizontal dashed line indicates uniform distribution. P-values were calculated using a two-sided paired *t* test. Place cells overrepresent rewarded locations at the end of the track in R1 and R2 which disappears in UR1 and UR2.

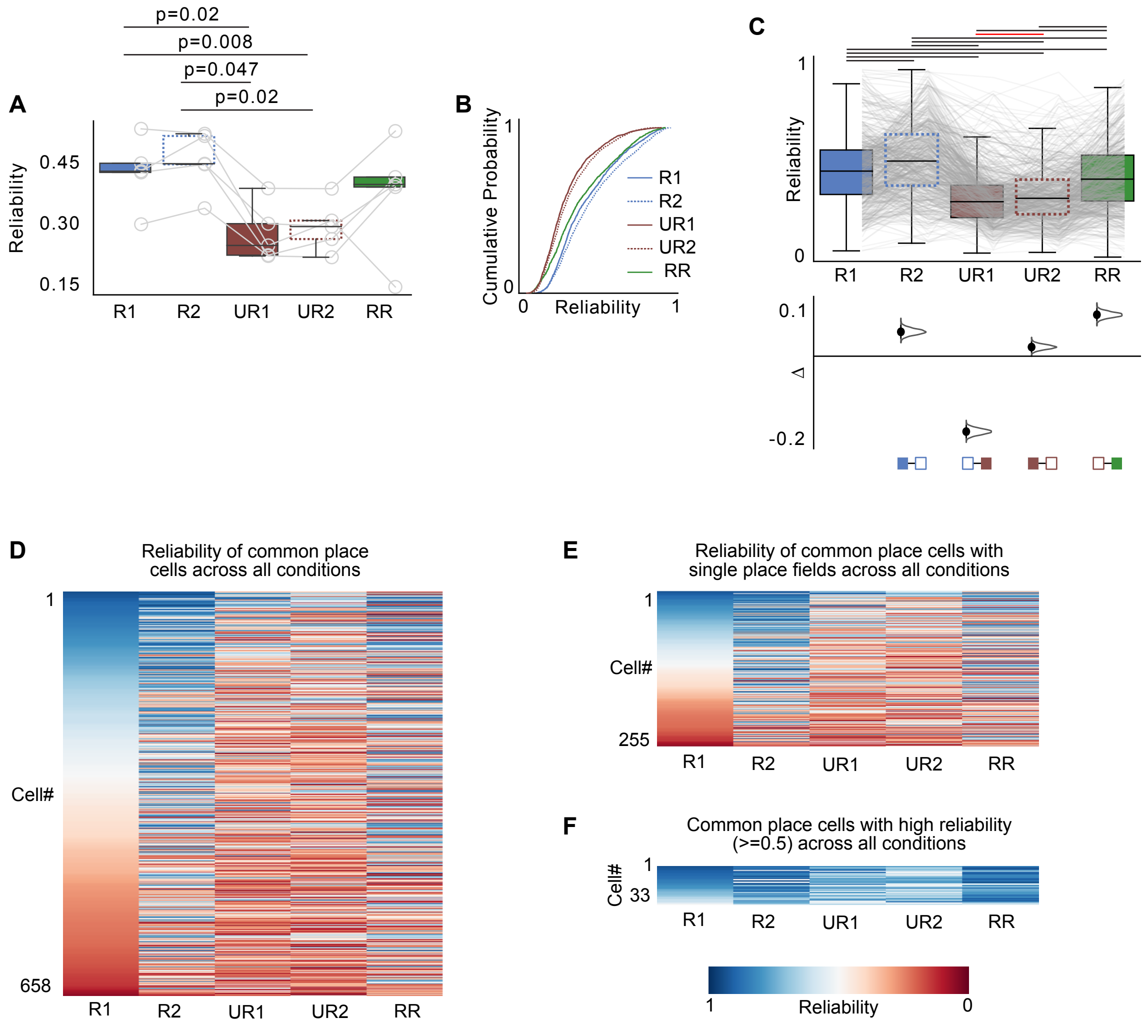


**Supplementary Figure 6. When reward expectation is low, place field reliability decreases. (A)** Boxplot of average reliability per animal of place cells defined in each condition. Each circle is an animal. P-values were obtained using a two-sided paired *t* test (**B**) Reliability of all place cells defined in each condition as a cumulative histogram. All relationships are significant **(C)** (Top) Boxplot of reliability of all common place cells. Place cells were defined in R1 and considered common if they had place fields in all other conditions. Grey lines indicate individual cells. Significant relationships are indicated by horizontal black lines on top (two-sided paired *t* test, $p<0.05$) and insignificant relationships are indicated in red. (Bottom) Bootstrapped mean differences ($\Delta$) with 95% CI (error bar). *X*-axis indicates the comparisons made. Bootstrapped comparisons were made between consecutive conditions. (D) Heatmaps show a different way of visualizing changes in reliability of place cells in each condition. Colormap is centered at 0.5, blue hues represent cells with high reliability (>0.5) and red hues represent cells with low reliability (<0.5). All common place cells are displayed in D, common place cells that retain single fields across all conditions are displayed in E and common place cells that maintain reliability of >=0.5 across all conditions are displayed in F. For full details on statistics and p-values, see **supplementary file 1**.

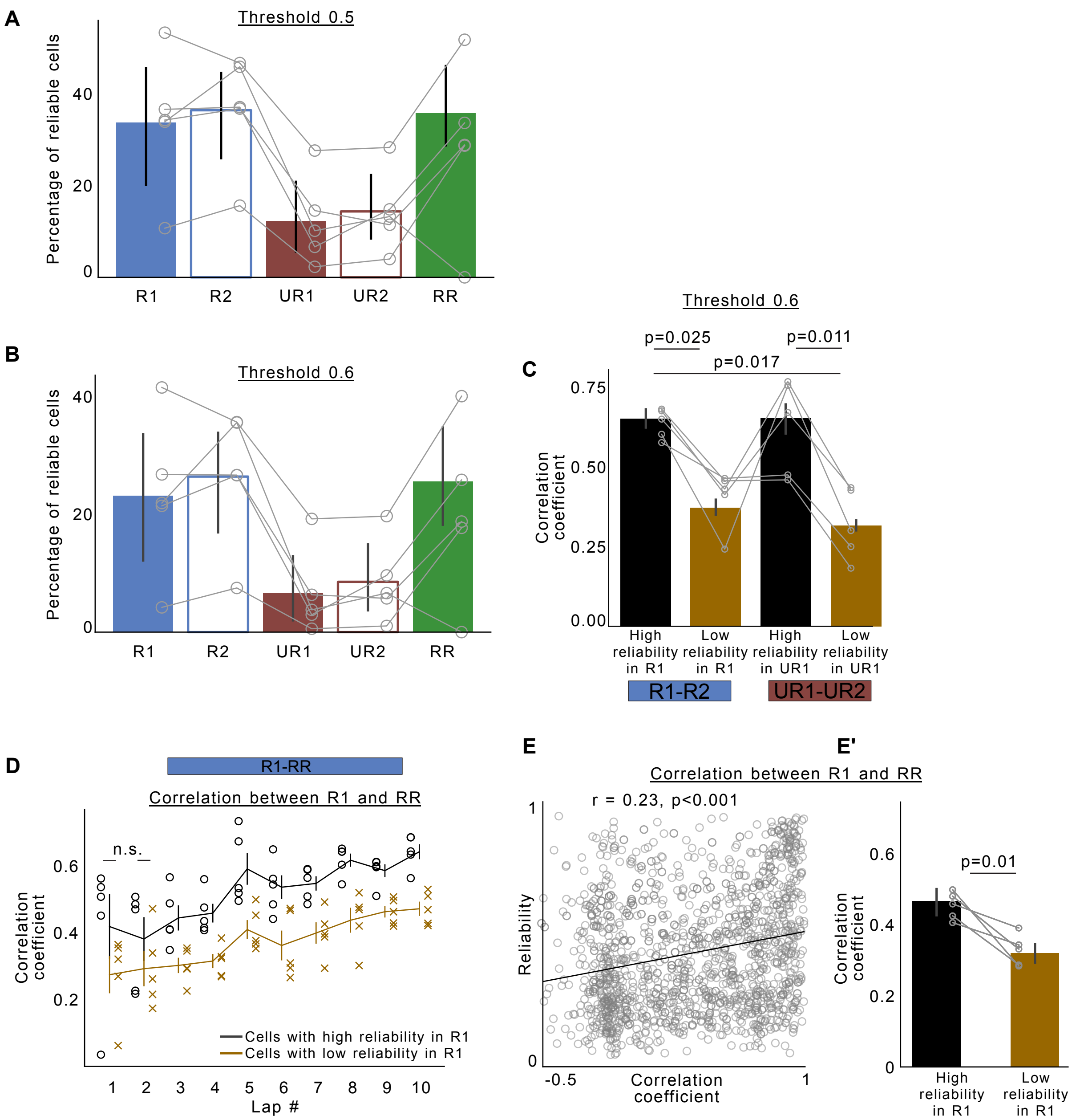


**Supplementary Figure 7. Varying threshold to define highly reliable place fields and place cells with more reliable fields from lap-to-lap are more stable across days between R1 and RR. (A-B)** Percentage of reliable cells calculated by dividing the number of reliable cells by the total number of place cells in each condition. Each circle is an individual animal. High and low reliable place fields were classified using a threshold of (**A**) 0.5 or (**B**) 0.6. (**C**) Animal wise averages of correlation coefficient between mean place fields on previous day (R1/UR1) with next day (R2/UR2) as indicated divided by high vs less reliable place fields obtained using a threshold of 0.6. P-values were obtained using a two-sided paired *t* test and corrected for multiple comparisons. **(D)** Average lap-wise correlation comparing last lap in R1 with first ten laps in RR. Place cells were defined in R1 and divided based on their reliability in R1. Cells with high reliability have reliability >=0.5 and cells with low reliability have reliability < 0.5. Circles and crosses are data from individual mice. Non-significant (n.s.) p-values are indicated, all others are significant. P-values were obtained using a two-sided paired *t* test between correlations of high reliable cells (black trace) vs low reliable cells (gold trace) **(E)** Scatter plot of place cell reliability in R1 (y-axis) compared to correlation between mean place fields in R1 and RR (x-axis). Each circle is a place cell defined in R1. Black line shows linear regression fit. r and p-values were obtained using Pearson's correlation coefficient. **(E')** Animal-wise averages of data in E divided by reliability using a threshold of 0.5. Circles indicate individual mice. P-values were obtained using a two-sided paired *t* test. For full details on statistics and p-values, see **supplementary file 1**. All error bars indicate 95% confidence interval

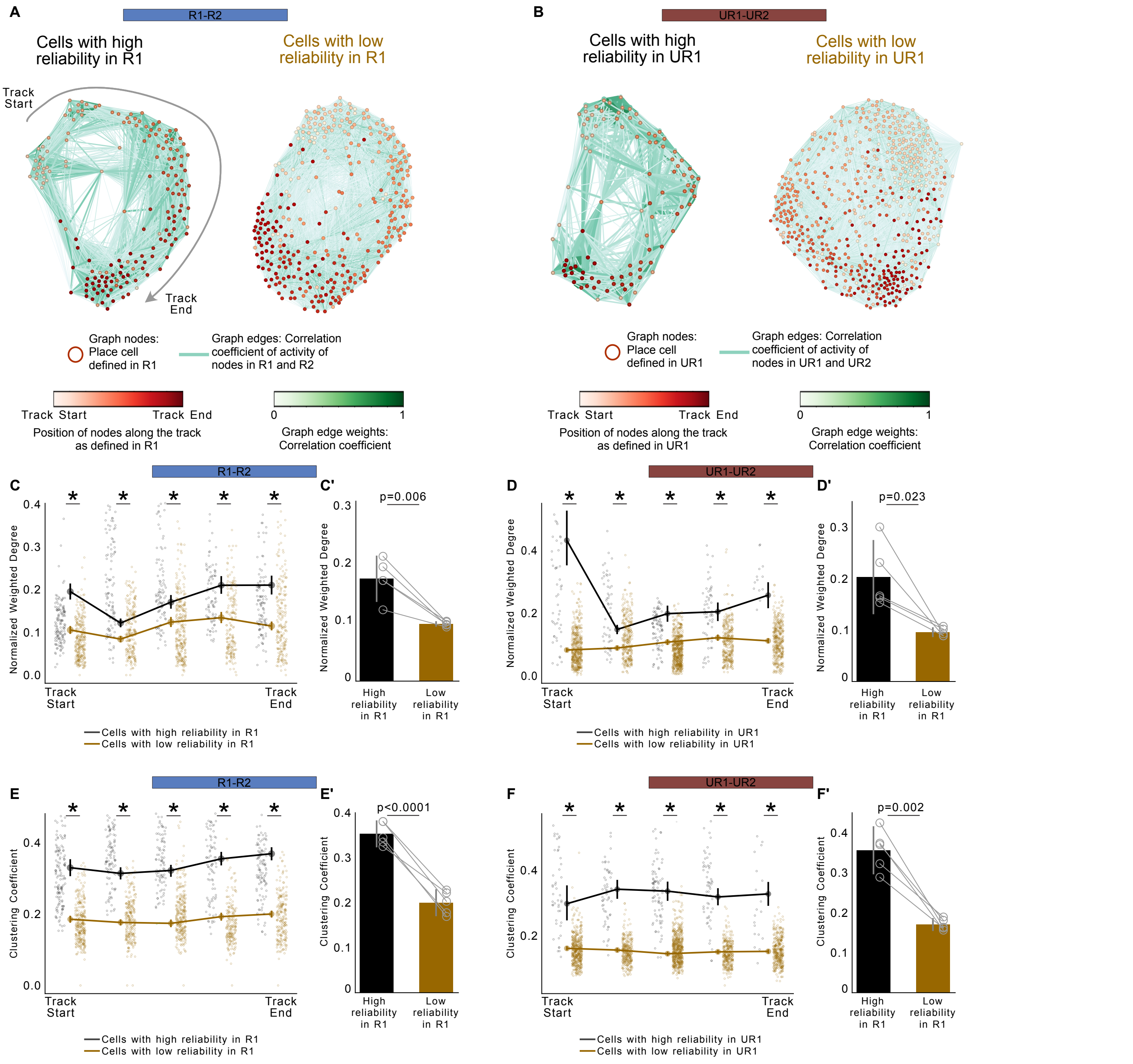


**Supplementary Figure 8. Cells with high lap-to-lap place field reliability exhibit stronger network correlation and connectivity across days. (A)** Example network graphs of neuronal co-activity. All graphs are from the same mouse. Each node (in shades of red) represents a place cell defined in R1 and divided by their reliability in R1. Edge weights (in shades of green) represent Pearson correlation coefficient between the activity of nodes in R1 with their activity in R2 (R-network). The graph is visualized using a Force Atlas 2 topology which arranges the nodes in a circular fashion grouping track positions that are closer to each other (see Methods). **(B)** Same as A but between UR1 and UR2 (UR-network). **(C)** Traces represent average weighted degrees from R-graphs of place cells with high (black) and low reliability (gold). Each dot is a node divided by its reliability with place field in that track location in R1 (**C'**) Average weighted degree plotted for each animal. (**D**) Same as C except for UR-network. (**E**) Average clustering coefficient of nodes binned by track position. Traces represent average and shading represents 95% Confidence Intervals. Each dot is a node divided by its reliability with place field in that track location in R1 (**E'**) Average clustering coefficient for each animal. (**F**) Same as E except for UR-networks. All error bars represent 95% confidence intervals. P-values were calculated using a Kolmogrov Smirnov (KS) test in C, D, E and F and a two-sided paired *t* test in C', D', E' and F'.

**Supplement File References:**

Krishnan, S., Heer, C., Cherian, C., & Sheffield, M. E. J. (2022). Reward expectation extinction restructures and degrades CA1 spatial maps through loss of a dopaminergic reward proximity signal. *Nature Communications*, *13*(1), 6662.

Stringer, C., Zhong, L., Syeda, A., Du, F., Kesa, M., & Pachitariu, M. (2023). Rastermap: a discovery method for neural population recordings. In *bioRxiv* (p. 2023.07.25.550571). https://doi.org/10.1101/2023.07.25.550571
